# No gender differences in predictive processing

**DOI:** 10.1101/2025.11.04.686317

**Authors:** Inés Botía, Bianka Brezóczki, Adrienn Holczer, Dezső Németh, Teodóra Vékony

**Affiliations:** Gran Canaria Cognitive Research Center, Atlántico Medio University, Las Palmas de Gran Canaria, Spain; Doctoral School of Psychology, ELTE Eötvös Loránd University, Budapest, Hungary; Institute of Psychology, ELTE Eötvös Loránd University, Budapest, Hungary; Brain, Memory and Language Research Group, Institute of Cognitive Neuroscience and Psychology, HUN-REN Research Centre for Natural Sciences, Budapest, Hungary; Centre de Recherche en Neurosciences de Lyon CRNL U1028 UMR5292, INSERM, CNRS, Université Claude Bernard Lyon 1, 69500 Bron, France; BML-NAP Research Group, Institute of Psychology, Eötvös Loránd University & Institute of Cognitive Neuroscience and Psychology, HUN-REN Research Centre for Natural Sciences, Budapest, Hungary

## Abstract

Statistical learning, defined as the implicit extraction of environmental regularities, is considered a fundamental cognitive mechanism that supports predictive processing across individuals, domains, and species. However, whether it is modulated by gender remains unclear. This study investigated potential gender-related differences in statistical learning using an age-matched sample of 129 women and 129 men (N = 258) who completed a well-established visuomotor probabilistic learning task. Statistical learning was revealed in both reaction time and accuracy measures, with participants responding faster and more accurately to high-probability than to low-probability trials. Critically, neither the magnitude nor the trajectory of statistical learning differed between women and men. Furthermore, no baseline differences in visuomotor performance interacted with learning metrics. Overall, these findings suggest that implicit statistical learning is a robust cognitive mechanism that is highly resilient to gender-related variation.

**Highlights:**

⌷ Implicit statistical learning was robust in both women and men, with no evidence for gender differences in the magnitude or trajectory of learning.
⌷ Although a minor baseline difference in general response speed was observed, this effect did not interact with or influence statistical learning.
⌷ Statistical learning was robust across menstrual cycle phases and hormonal contraceptive use groups.
⌷ The findings suggest that the "predictive brain" relies on fundamental, evolutionarily conserved mechanisms that are largely resilient to gender-based variation.

---

The human brain continuously builds predictive models of the world by extracting statistical regularities from sensory input, a fundamental capacity known as statistical learning (e.g., Aslin, 2017; Fiser & Aslin, 2002; Saffran et al., 1996). This implicit mechanism is considered a key element of cognition (Santolin & Saffran, 2018; Santolin et al., 2020) and plays a central role in domains ranging from language acquisition (Saffran et al., 1996) to skill development (Janacsek et al., 2012). Its ubiquity across domains has led to the prevailing view that it reflects a broadly shared and evolutionarily conserved mechanism (Clark, 2013; Frost et al., 2015; Siegelman & Frost, 2015). This assumption contrasts with the substantial variability observed across cognitive domains, particularly the persistent debate surrounding sex differences in brain structure and function (Giofrè et al., 2024; Wierenga et al., 2018). This debate can be framed within the gender similarities hypothesis, which proposes that males and females are largely alike across most psychological and cognitive domains (Hyde, 2005). Meta-analytic evidence shows that although substantial gender differences exist in some physical and motor abilities, such as sports performance and throwing ability (Thomas & French, 1985), differences in core cognitive skills are generally small or absent (Hyde, 2005). Given that statistical learning supports domains such as language (Isbilen & Christiansen, 2022; Ren et al., 2023) and reading (Seidenberg et al., 2018), it might be expected to follow a similar pattern of sex similarity. However, evidence from language development is mixed, with some studies reporting small female advantages (Lange et al., 2016) and others finding little or no evidence for sex differences once environmental factors are considered (Ibbotson & Browne, 2024; Rinaldi et al., 2023). Whether foundational predictive processes truly operate uniformly across genders, therefore, remains an open question. Answering it is critical not only for defining the fundamental properties of cognitive architecture but also for understanding the origins of gender-based differences observed in higher-order cognitive domains.

Differences in cognitive domains are often attributed to the influence of sex hormones, which exert both long-term organizational effects on brain development (i.e., shaping neural structure and circuitry during development) and short-term activational effects on adult neural function (i.e., transiently modulating cognition and behavior through fluctuations in hormone levels) (Cahill, 2006). Nonetheless, this body of research has concentrated on explicit cognitive tasks requiring conscious reasoning or simple deterministic sequences (Höbler et al., 2022). In the domain of statistical processing, previous studies have employed diverse explicit tasks, often yielding inconsistent findings regarding gender differences. Such inconsistencies may partly arise because performance in these tasks is influenced not only by statistical processing itself but also by individual differences in formal experience, cognitive ability, and thinking dispositions (Martin et al., 2017), as well as working memory capacity and conscious segmentation strategies. Furthermore, many foundational studies on motor and sequence learning either relied exclusively on male participants or have not yet comprehensively examined gender-based differences, compounding the inconsistencies in the literature (Höbler et al., 2022). This leaves a critical question unanswered: do gender-related factors extend to the implicit, largely automatic mechanisms that underpin predictive cognition? Since statistical learning operates largely outside of conscious awareness, it provides a crucial test case for the reach of gender-based influences. By employing an implicit, probabilistic paradigm, such as the Alternating Serial Reaction Time task (ASRT), it is possible to bypass the conscious strategies that confound explicit tasks, ensuring that statistical regularities are extracted without participants developing explicit awareness of the hidden sequence (Höbler et al., 2022; Vékony et al., 2021). Determining whether this fundamental mechanism is influenced by gender factors is crucial for understanding the consistency of cognitive architecture across individuals and the origins of variation in the brain’s predictive processes.

The lack of investigation into these factors is particularly striking given that hormonal variables known to differ between the genders, such as menstrual cycle phase (Milad et al., 2010) and hormonal contraceptive use, may interfere with or modulate performance on learning and memory tasks (Milad et al., 2010). While sex differences have also been reported in more general domains, such as motor reaction time, findings are often inconsistent and task-dependent (e.g., Deary & Der, 2005; but see Bunce et al., 2008; Kalb et al., 2004). Despite these extensive lines of inquiry, the role of gender in statistical learning and predictive processing remains largely unexamined. The majority of studies either do not analyze for gender differences or use mixed-gender samples without specific examination, leaving a critical blind spot in the literature (Maki et al., 2002). The significance of this oversight is underscored by hints from neuroimaging and clinical studies of sex-specific patterns in statistical learning and the vulnerability of its underlying neural substrates (Colla et al., 2003).

Here, we investigated gender-based variation in implicit statistical learning using a well-established visuomotor statistical learning task (Farkas et al., 2024) in a cohort of men and women participants. The task allowed us to dissociate distinct cognitive components, including general reaction time, visuomotor learning (a speed-up independent of statistical regularities), and both the magnitude and trajectory of statistical learning. In addition to the primary gender comparison, we conducted extended analyses on hormonal contraceptive use and menstrual cycle phase, which are reported in the Supplementary Materials. This investigation provides a comprehensive examination of gender differences in this fundamental cognitive mechanism, offering crucial insights into the universality of the brain’s ability to learn environmental patterns.

## Method

### Participants

A total of 596 healthy young individuals participated in an online study. To ensure good data quality, only those who met all inclusion criteria were retained for analysis. A total of 123 participants were excluded for the following reasons: 14 participants failed the attention check questions designed to ensure task compliance, 14 did not complete the full experiment, 2 restarted the task, 60 self-reported a psychiatric diagnosis, 22 failed to meet the accuracy criterion (> 80%), and 11 reported taking medication that affects the central nervous system. Of the excluded participants, 89 identified as women and 31 as men (1 selected “Other,” and 2 selected “Prefer not to answer.”), corresponding to exclusion rates of 21.0% and 19.4% of the initially recruited women and men samples, respectively.

After applying exclusion criteria, the final sample comprised 473 participants. Gender identity was assessed through self-report. Of the final sample, 334 participants identified as women, 129 as men (M_a*ge*_ = 22 years, SD*_Age_* = 4.51), and 10 participants either identified with another gender, preferred not to disclose their gender, or did not provide a response. Given the unequal group sizes, an age-matched female subsample of 129 women (M_age_ = 22 years, SD*_Age_* = 4.51) was constructed using greedy nearest-neighbor matching (1:1, without replacement) implemented in R with the *dplyr* package (*tidyverse*) and matched to the male group on age for between-gender comparisons. Additionally, exploratory within-women analyses were conducted to examine potential hormonal influences on visuomotor performance and statistical learning. These analyses compared women using hormonal contraceptives with age-matched non-users and examined differences across self-reported menstrual cycle phases among naturally cycling women. Detailed information regarding participant selection and subgroup characteristics is provided in the Supplementary Materials.

The present study constitutes a secondary analysis of an existing dataset that was originally collected as part of a broader research project investigating statistical learning. Some data from this dataset have been published previously in studies addressing different research questions and using partial samples; however, none examined the full dataset or addressed the research questions investigated in the present study (Brezoczki et al., 2025a; Brezoczki et al., 2025b; Farkas et al., 2026; Hann et al., 2025; Nagy et al., 2025).

All participants provided informed consent prior to enrollment. Participants received course credits for taking part in the study. The study was approved by the Research Ethics Committee of Eötvös Loránd University, Budapest, Hungary (protocol number: 2021/504), and conducted in accordance with the principles of the Declaration of Helsinki.

### Task

#### Alternating Serial Reaction Time task (ASRT)

To measure implicit statistical learning, we employed the ASRT task, a visuomotor learning paradigm with well-established reliability (Farkas et al., 2024). The task was programmed in JavaScript using the jsPsych library (De Leeuw, 2015) and consisted of 15 blocks of 80 trials each, preceded by 2 practice blocks.

On each trial, a stimulus, a drawing of a dog’s head, appeared in one of four horizontally arranged positions on the screen. Participants were instructed to respond as quickly and accurately as possible by pressing the corresponding key (“S”, “F”, “J”, or “L”, from left to right). Participants used both hands, the left middle finger for the “S” key, the left index finger for the “F” key, the right index finger for the “J” key, and the right middle finger for the “L” key. Participants maintained this fixed finger-to-key mapping throughout the task. The stimulus remained visible until the correct response was provided, followed by a 120 ms response-to-stimulus interval (Figure 1).

**Figure 1.**
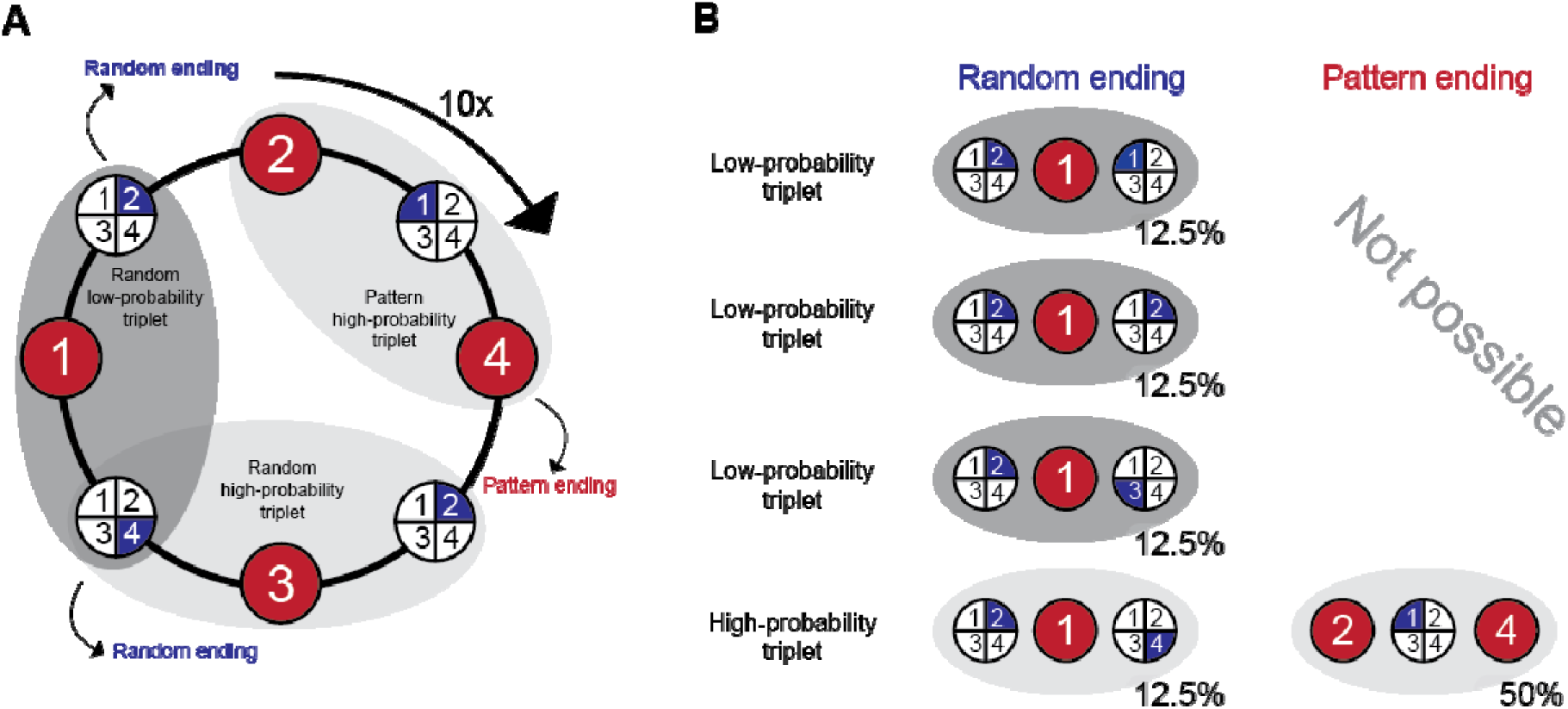
Structure of the ASRT task. **A.** The task follows an eight-element statistical sequence in which fixed pattern positions (red) alternate with random positions (blue), forming series such as 2–r–4–r–3–r–1–r. This alternating structure generates “triplets” (three consecutive trials), which can end either on a pattern element (pattern ending) or on a random element (random ending). **B.** As a result, some triplets occur more frequently (high-probability triplets) than others (low-probability triplets). High-probability triplets can occur in two different ways: either the first and third elements belong to the fixed pattern sequence (P–r–P structure) or the middle element belongs to the fixed pattern sequence (r–P–r structure). In contrast, low-probability triplets can only occur in r–P–r structures because their final element cannot appear as part of the fixed pattern sequence.

Participants were not aware that the stimuli followed an alternating eight-element sequence composed of fixed pattern (P) and random (r) positions (e.g., 2–r–4–r–3–r–1–r). This alternating structure generated “triplets,” defined as runs of three consecutive trials. Because the pattern elements followed a predetermined order, some triplets occurred more frequently than others. Specifically, high-probability triplets could occur in both P–r–P and r–P–r structures, whereas low-probability triplets could only occur in r–P–r structures (Figure 1). For example, triplets like 2–X–4 or 3–X–1 (where X is any middle element) occurred more frequently, since their final element often appeared as part of the fixed pattern (and, in some cases, as a random element as well). In contrast, triplets such as 2–X– 1 or 2–X–3 could only occur with lower probability because their final element could only appear in random positions. Importantly, trials were classified according to whether they constituted the final element of a high- or low-probability triplet (Figure 1). In total, 62.5% of the trials were the last element of a high-probability triplet (high-probability trials), while the remaining 37.5% formed low-probability triplets (low-probability trials).

### Procedure

The study was administered online using the Gorilla Experiment Builder platform (https://www.gorilla.sc) (Anwyl-Irvine et al., 2020). Participants first read an introductory briefing and provided informed consent. Then, they completed the ASRT task, which started with two practice blocks composed entirely of random sequences. These practice blocks were excluded from subsequent analyses. After the practice phase, participants completed 15 experimental blocks. At the end of each block, participants received feedback on their mean reaction time and accuracy. After completing a block, participants initiated the next one by pressing a key, allowing them to proceed at their own pace.

Following the ASRT task, participants answered a series of demographic questions covering topics such as socioeconomic background, sleep patterns, general health, handedness, menstrual cycle, and hormonal contraceptive use. Due to the remote nature of the study, participants were also asked to provide brief information about the conditions under which they completed the task.

Participants were not informed about the underlying sequence and were later asked whether they had noticed any hidden patterns, with a request to elaborate if so. None were able to accurately identify the alternating sequence, confirming that learning remained implicit, consistent with previous findings (Horváth et al., 2022; Vékony et al., 2022).

### Statistical analysis

We used R (version 4.4.3) to process behavioral data from the ASRT task. Each trial was categorized based on the two preceding trials as the last element of a high- or low-probability triplet. Following standard practice (e.g., Farkas et al., 2024; Vékony et al., 2025), trills (e.g., 1-2-1, 2-1-2, 3-1-3, 1-4-1), repetitions (e.g., 2-2-2), the first two trials of each block, and trials with RT below 100 ms or above 1,000 ms were excluded from the analyses. Trials exceeding 3 median absolute deviations from the individual median were also removed. Inaccurate responses were also removed from the RT analyses.

To investigate the trial-by-trial dynamics of statistical learning, both RTs and accuracy were analyzed using Generalized Linear Mixed Models (GLMMs) implemented via the *afex* package in R. Because RT distributions are typically right-skewed, the RT model was fitted using an inverse Gaussian distribution with a log link function, which accommodates the characteristics of continuous RT data without requiring prior mathematical transformations. Accuracy was analyzed using a binomial GLMM with a logit link function. Both models included epoch (Epoch 1 vs. Epoch 2 vs. Epoch 3) and triplet type (high-probability vs. low-probability) as within-subject fixed effects, and gender (female vs. male) as a between-subject fixed effect, along with all possible interaction terms. Epoch was treated as an ordered categorical factor, and global contrast options were set to sum-to-zero for unordered factors and polynomial contrasts for ordered factors. We specified a random effects structure that included random intercepts for participants, as well as by-participant random slopes for both epoch and triplet type (representing the maximal converging random effects structure). The model optimizer was set to allow a maximum of 1,000,000 function evaluations to ensure convergence. The statistical significance of the fixed effects was evaluated using Likelihood Ratio Tests (LRT) by comparing the full model against sequentially reduced models. The alpha level was set at .05, and p-values between .05 and .10 were interpreted as trends.

To move beyond traditional null-hypothesis significance testing and quantify the evidence for practical equivalence, a quasi-Bayesian Region of Practical Equivalence (ROPE) analysis was conducted. This approach utilizes a multivariate normal (MVN) approximation of the contrast estimates derived from the GLMMs. Using the *emmeans* and *MASS* packages, we simulated the posterior distribution of the estimated marginal means by drawing 40,000 samples from the multivariate normal distribution parameterized by the models’ fixed-effect estimates and their associated variance-covariance matrices. We defined ROPEs a priori for both outcomes: for RT, a ±1% change on the multiplicative (log-link) scale; for accuracy, a ±1% difference around p = .90 (converted to the log-odds scale for the binomial GLMM). For each contrast of interest, we calculated the median percentage change, the 95% Credible Intervals (CrI), and the exact percentage of the simulated posterior distribution that fell strictly within the predefined ROPE. Following standard decision rules, if more than 95% of the posterior distribution fell within the ROPE, it was considered strong evidence that the effect size is practically equivalent to zero, allowing us to statistically support the null hypothesis.

All data and code required to reproduce the results of this study are available on the OSF repository (https://osf.io/u82w9/overview).

## Results

### Lack of gender differences in statistical learning - RT measures

To analyze the trial-by-trial dynamics of statistical learning, a GLMM was fitted to the RTs and accuracies. The RT model confirmed robust visuomotor learning throughout the task, evidenced by a highly significant main effect of epoch (□^2^(2) = 43.96, *p* < .001) (Figure 2A). Furthermore, statistical learning was clearly established, as the main effect of triplet type was highly significant (□^2^(1) = 56.81, *p* < .001), indicating that participants responded faster to high-probability versus low-probability triplets (Figure 3A). The model also revealed a significant interaction between epoch and triplet type (□^2^(2) = 26.89, *p* < .001), demonstrating that the magnitude of statistical learning evolved across the epochs (Figure 3A).

**Figure 2.**
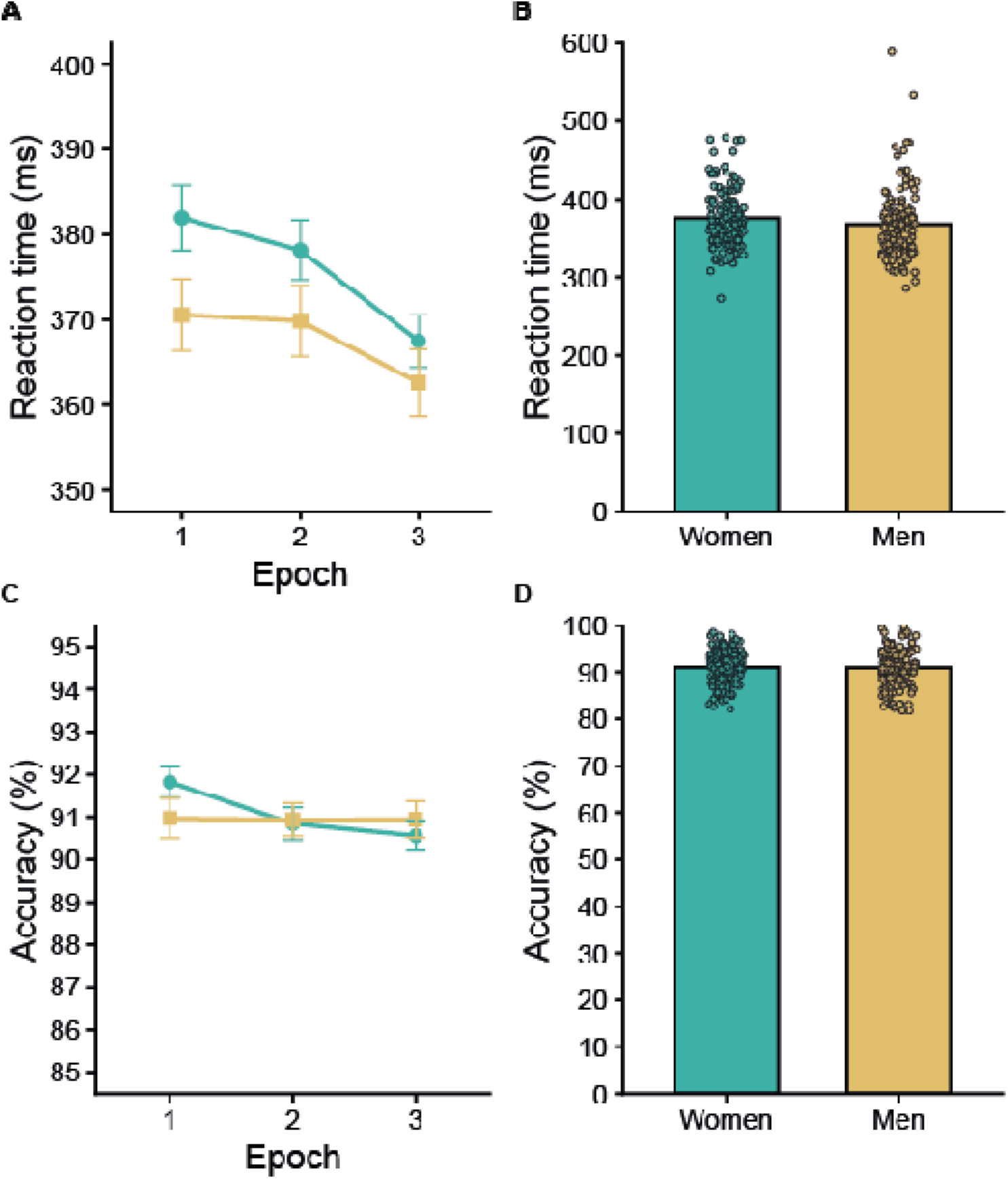
Lack of gender differences in visuomotor learning. Women are represented in green and men in yellow throughout all panels. In Panels A and C, lines represent group means, and error bars indicate the standard error of the mean (SEM). In Panels B and D, bars represent group means, and individual dots represent participant-level scores. **A.** Reaction times (RTs; ms) across the three epochs of the task. RTs decreased over time, indicating visuomotor learning. Women exhibited somewhat slower RTs than men (ROPE analysis); however, the pattern of improvement across epochs was comparable between groups. **B.** Overall reaction time, computed as the mean of Epochs 1–3. Women exhibited somewhat slower reaction times than men, consistent with a baseline processing-speed difference. Importantly, this difference was unrelated to predictive processing and did not influence the magnitude or trajectory of statistical learning. **C.** Accuracy (%) across the three epochs of the task, averaged across triplet types. The pattern of accuracy across epochs was highly similar in women and men, providing no evidence for gender-related differences in learning. **D.** Overall accuracy, computed as the mean of Epochs 1–3. Women and men showed comparable levels of accuracy, consistent with the absence of significant gender effects in the accuracy model.

**Figure 3.**
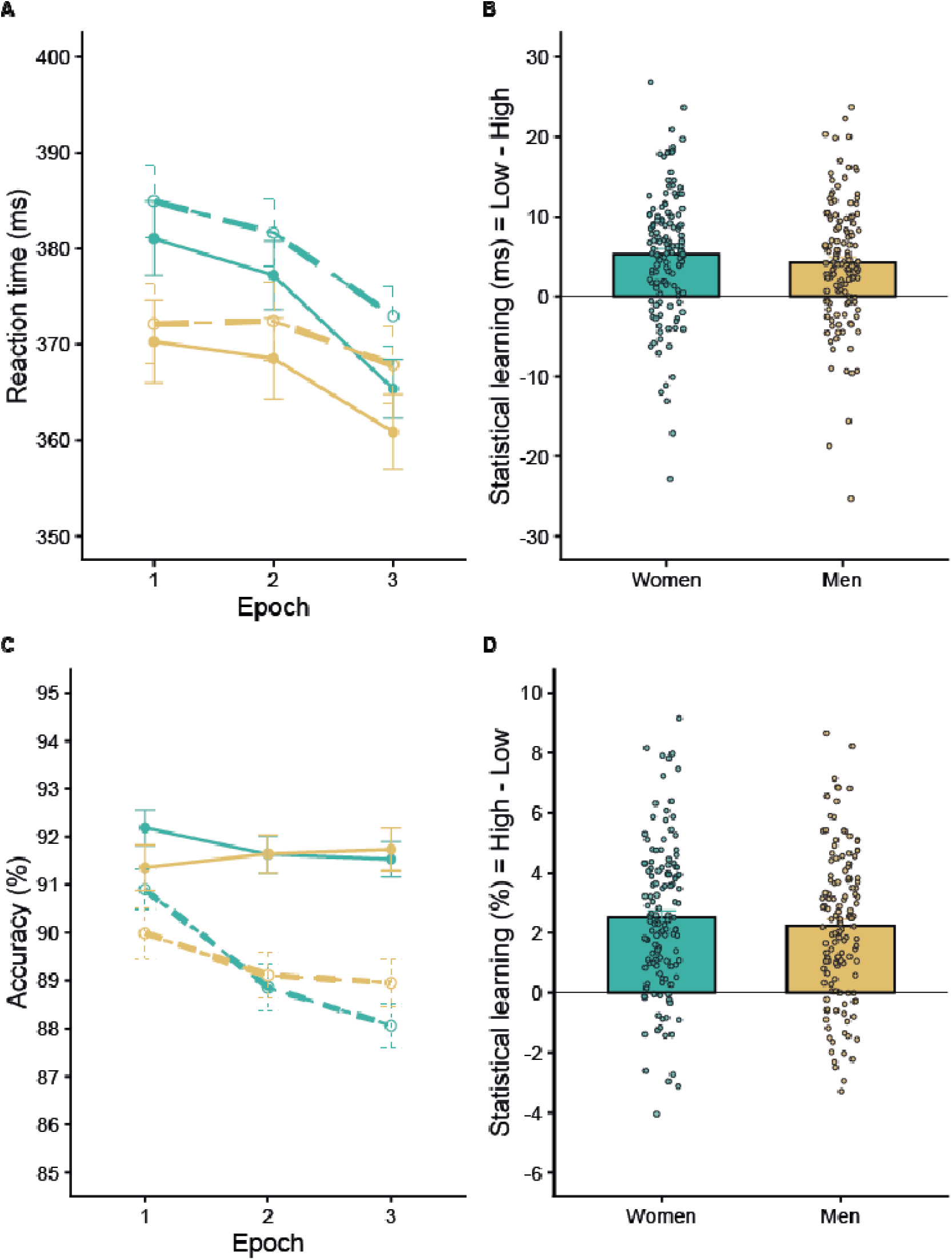
Absence of gender effects in statistical learning. Women are represented in green and men in yellow throughout all panels. In Panels A and C, lines represent group means, and error bars indicate the standard error of the mean (SEM). Solid lines represent high-probability triplets and dashed lines represent low-probability triplets. In Panels B and D, bars represent group means, and individual dots represent participant-level statistical learning indices. **A.** Reaction times (RTs; ms) across the three epochs for high-probability (solid lines) and low-probability (dashed lines) triplets. Participants responded faster to high-probability than to low-probability triplets, indicating robust statistical learning. This learning effect evolved across epochs but did not differ between women and men. **B.** Statistical learning in RT, computed as Low − High (ms) averaged across Epochs 1–3. Statistical learning was evident in both groups and was comparable between women and men, consistent with the absence of gender differences in sensitivity to statistical regularities. **C.** Accuracy (%) across the three epochs for high-probability (solid lines) and low-probability (dashed lines) triplets. Participants were more accurate on high-probability than on low-probability triplets, further demonstrating robust statistical learning. We found no reliable gender differences. **D.** Statistical learning in accuracy, computed as High − Low (%) averaged across Epochs 1–3. Statistical learning was evident in both groups and showed no meaningful differences between women and men, consistent with the strong evidence for practical equivalence observed in the ROPE analysis.

Addressing the primary research question regarding gender differences, the GLMM did not provide evidence for a main effect of gender (□^2^(1) = 1.54, *p* = .214) or of an interaction between gender and epoch (□^2^(2) = 2.60, *p* = .272). Most importantly, the interaction between gender and triplet type was not significant (□^2^(1) = 0.88, *p* = .348), indicating that women and men did not differ in their sensitivity to statistical regularities. Additionally, the three-way interaction between gender, epoch, and triplet type was absent (□^2^(2) = 0.27, *p* = .874), confirming that the trajectory of statistical learning over time was statistically indistinguishable between the two genders (Figure 3B).

To move beyond traditional null-hypothesis significance testing and quantify the evidence for equivalence between genders, a Quasi-Bayesian (ROPE) analysis was conducted on the GLMM contrasts. Visuomotor learning was further validated, showing a robust median RT decrease of -2.62% between the third and first epochs (95% CrI [-3.01%, -2.23%]) (Figure 2A), with 0% of the posterior distribution falling within the ROPE. Similarly, the overall statistical learning effect showed a median difference of 1.26% (95% CrI [1.04%, 1.48%]), with only 0.98% of the distribution inside the ROPE. The temporal dynamics of this learning were captured by the epoch and triplet interaction (Epoch 3 minus Epoch 1), which yielded a median difference of 1.04% (95% CrI [0.63%, 1.44%], 43.36% in ROPE).

Crucially, the Quasi-Bayesian analysis provided overwhelming evidence for the absence of gender differences in statistical learning. The critical interaction between gender and triplet type, averaged across all epochs, had a median estimate of just 0.27% (95% CrI [-0.15%, 0.69%]) (Figure 3B). Strikingly, 99.97% of this posterior distribution fell strictly within the ROPE, offering decisive support for practical equivalence between women and men in statistical learning. The three-way interactions comparing the gender gap in statistical learning between specific epochs are also firmly localized within the ROPE, such as the Epoch 3 minus Epoch 1 contrast (97.45% in ROPE) and the Epoch 3 minus Epoch 2 contrast (98.36% in ROPE). As a secondary finding, the analysis detected a baseline difference in general RTs, with women being somewhat slower than men (Median difference = 2.47%, 95% CrI [1.69%, 3.26%], 0% in ROPE). However, this baseline processing speed difference did not interact with the predictive processing mechanisms, solidifying the conclusion that statistical learning functions equivalently across genders.

### Lack of gender differences in statistical learning - Accuracy measures

A GLMM was fitted to the accuracies as well, with the same fixed and random effect structure. The accuracy model confirmed robust visuomotor learning throughout the task, evidenced by a highly significant main effect of epoch (□^2^(2) = 21.48, *p* < .001) (Figure 2C). Furthermore, statistical learning was clearly established, as the main effect of triplet type was highly significant (□^2^(1) = 176.02, *p* < .001), indicating that participants responded more accurately to high-probability versus low-probability triplets. The model also revealed a significant interaction between epoch and triplet type (□^2^(2) = 28.69, *p* < .001), demonstrating that the magnitude of statistical learning evolved across the epochs (Figure 3C).

The GLMM provided no evidence for a main effect of gender (□^2^(1) = 0.13, *p* = .716), but a trend toward an interaction between gender and epoch (□^2^(2) = 4.70, *p* = .096). Most importantly, the interaction between gender and triplet type was not significant (□^2^(1) = 0.30, *p* = .582), indicating that males and females did not differ in statistical learning. Additionally, the three-way interaction between gender, epoch, and triplet type was completely absent (□^2^(2) = 0.43, *p* = .808), confirming that the trajectory of statistical learning over time was similar between the two genders (Figure 3D).

We conducted a ROPE analysis on binomial-GLMM contrasts, using a pre-defined ROPE corresponding to ±1 percentage point around p = .90 (|log-odds| <= 0.1164). The analysis confirmed robust learning effects in accuracy: the overall statistical learning contrast showed a median difference of -3.07% (95% CrI [-3.49%, -2.67%]), with 0% of the posterior inside the ROPE. Consistently, temporal changes in learning were also substantial, as seen in the epoch by triplet interactions for Epoch 3 minus Epoch 1 (Median = -1.84%, 95% CrI [-2.61%, -1.12%], 2.12% in ROPE) and Epoch 2 minus Epoch 1 (Median = -1.33%, 95% CrI [-2.07%, -0.63%], 25.72% in ROPE).

The critical gender by triplet interaction, averaged across epochs, had a median estimate of 0.18% (95% CrI [-0.47%, 0.80%]), with 99.53% of the posterior distribution securely inside the ROPE. Epoch-specific gender by triplet contrasts similarly concentrated near practical equivalence (Epoch 1: 95.86% in ROPE; Epoch 2: 95.12%; Epoch 3: 91.21%). Three-way gender by epoch by triplet contrasts also showed high ROPE inclusion (Epoch 2 minus 1: 86.59%; Epoch 3 minus 1: 82.00%; Epoch 3 minus 2: 88.76%), indicating that the temporal trajectories of statistical learning do not differ by gender. Finally, other temporal and baseline contrasts largely supported equivalence or lacked robust deviations, with the gender by epoch interaction (Epoch 3 minus 1) showing the widest spread, though still maintaining a median close to the equivalence bounds (Median = 1.28%, 95% CrI [0.12%, 2.31%], 31.13% in ROPE).

## Discussion

A central question in cognitive science is whether fundamental learning mechanisms are universal or subject to individual variation, such as gender. While sex-based differences have been observed in higher-order cognitive domains like language and spatial ability (Halpern, 2000; Miller & Halpern, 2014), it remains unclear whether these extend to more foundational processes, such as predictive processing. To address this issue, we analyzed data from 473 young adults who completed the ASRT task in an online setting. Using a large sample and a well-established online ASRT paradigm, we constructed an age-matched sample of 129 women and 129 men (N = 258) for the gender-comparison analyses and compared men and women on multiple components of visuomotor performance and statistical learning. Our results provide a clear picture: statistical learning was robust across participants, and we found no evidence that it is modulated by gender. Our findings align with previous evidence showing that statistical learning functions as a broad and flexible mechanism, observable across sensory modalities, tasks, and species (Armstrong et al., 2017; Sonnweber et al., 2015). Although men initially responded slightly faster than women, this difference did not influence overall learning performance. Overall, statistical learning appeared to be stable and comparable between genders, suggesting that this form of implicit learning is largely unaffected by the biological factors that often differentiate male and female cognition in other domains (Becker et al., 2016; Kimura & Hampson, 1994).

The absence of gender-related differences in statistical learning aligns with the gender similarities hypothesis, which emphasizes that males and females tend to be more alike than different across most psychological variables (Hyde, 2016). Earlier reviews reached similar conclusions, noting that although some exceptions exist, cognitive and behavioral traits generally show more overlap than divergence between the sexes (Hyde, 2005). We propose that the resilience of statistical learning to gender-based variation stems from its automatic nature. Unlike functions that recruit executive control and are susceptible to strategic modulation, statistical learning is thought to operate largely independently of conscious oversight (Aslin & Newport, 2012; Frost et al., 2015; Saffran et al., 1996; Vékony et al., 2021). Supporting this view, comparative studies report similar outcomes across species: in chicks, males and females showed comparable preferences for statistical regularities (Santolin et al., 2016), and in rodents, both sexes learned a sequence memory task at similar rates, with only minor strategic differences (e.g., shorter poke times in males) that did not affect overall accuracy; once the task was fully acquired, no sex differences remained (Jayachandran et al., 2022). Together, these observations reinforce the notion that statistical learning is a fundamental and automatic cognitive mechanism that operates similarly across genders.

The formal likelihood ratio test of the gender main effect was non-significant, whereas the marginal estimates from the ROPE analysis pointed to a small baseline difference in general reaction times, with women responding somewhat slower than men. We treat this baseline difference as a secondary, descriptive observation rather than a primary effect. The two analyses address different quantities: the likelihood ratio test evaluates the overall contribution of the gender term to the model, while the marginal contrast isolates the estimated difference in general response speed, so they are complementary rather than contradictory. Numerous studies have reported that men tend to respond faster than women in both simple and choice RT tasks, a trend observed from childhood (Goodenough, 1935) into adulthood (Christensen et al., 2001; Fairweather & Uttal, 1972; Reed et al., 2004), although some studies found no differences (Bunce et al., 2008; Kalb et al., 2004). Large-scale investigations confirm that women’s RTs are often slower and more variable, with the magnitude of the difference shifting across the lifespan (Deary & Der, 2005). More fine-grained analyses suggest that women are faster in decision time, while men are faster in movement time, leading to relatively stable overall RT between genders (Hodgkins, 1963; Fairweather & Uttal, 1972; Landauer et al., 1980). This pattern is consistent with evidence that sex-related differences in motor performance may stem from peripheral factors, such as aerobic capacity or muscle mass (Rezende et al., 2006), as well as central nervous system and developmental contributions (Davies & Rose, 2000). Moreover, after puberty, males may benefit more from identical motor training experiences in terms of procedural learning (Dorfberger et al., 2009). Overall, these patterns suggest that observed gender differences reflect general motor performance factors rather than differences in statistical learning mechanisms.

To assess potential within-gender hormonal modulation of visuomotor performance and statistical learning, we additionally conducted exploratory subgroup analyses in women. Specifically, we compared users and non-users of hormonal contraceptives (n = 70 per group), and we examined performance across menstrual cycle phases in a subsample of naturally cycling women (n = 251). Full methodological details and complete statistical outputs are provided in the Supplementary Methods (Participants and Statistical Analysis sections); a brief interpretative overview is included here to contextualize the primary findings. Although there is substantial evidence that estradiol, progesterone, and hormonal contraceptives can modulate certain forms of learning (Milad et al., 2010; Zeidan et al., 2011), studies directly comparing women who use hormonal contraceptives and those with natural menstrual cycles remain scarce. Across both RT and accuracy measures, hormonal contraceptive use was unrelated to baseline performance, overall task improvement, and statistical learning (see Supplementary Results: "No evidence that hormonal contraceptive use influences visuomotor or statistical learning in reaction time measures" and "No evidence that hormonal contraceptive use influences statistical learning in accuracy measures"). Importantly, contraceptive users and non-users showed comparable sensitivity to the probabilistic structure of the task, and the trajectory of learning across epochs was virtually identical in the two groups. These findings extend previous reports suggesting that some forms of implicit or probabilistic learning may be relatively robust to hormonal status (Maki et al., 2002), even though other cognitive domains appear sensitive to endocrine fluctuations. Notably, while serotonergic pathways show sensitivity to hormonal status (Moses et al., 2000), our findings suggest that predictive processing mechanisms supporting statistical learning remain largely preserved regardless of contraceptive use.

The literature on menstrual cycle effects in cognition has produced inconsistent results (Solís-Ortiz & Corsi-Cabrera, 2008). Some studies report phase-related changes in attention or motor performance (Álvarez-San Millán et al., 2021; Hjelmervik et al., 2014), as well as in motor coordination and perceptual-spatial skills, with improved motor speed but impaired spatial performance during the midluteal phase relative to menses (Hampson & Kimura, 1988), but others did not replicate these effects (Pletzer & Harris, 2018). Clinical evidence also shows that estrogen receptor modulation can enhance probabilistic association learning in schizophrenia (Kindler et al., 2015). Consistent with the absence of robust phase effects, studies in rodents have also reported no differences between phases of the estrous cycle in sequence memory (Jayachandran et al., 2022). We found no reliable evidence that the menstrual cycle phase modulated statistical learning. Neither RT-nor accuracy-based learning differed across phases, and the temporal dynamics of learning were comparable throughout the task (see Supplementary Results: "No evidence that menstrual cycle phase influences statistical learning in reaction time measures" and "No evidence that menstrual cycle phase influences statistical learning in accuracy measures"). Taken together, the findings provide little support for the view that naturally occurring hormonal fluctuations substantially influence predictive processing as measured by the ASRT task.

Several avenues for future research can build on the strengths of this study and refine the current approach. The fully online nature of the study, while advantageous for recruiting a large sample, limited experimental control over factors such as hardware variability, internet latency, and environmental distractions, potentially introducing noise into behavioral measures.

Overall, our findings demonstrate that implicit statistical learning is a robust and stable cognitive mechanism, remaining unaffected by gender, hormonal contraceptive use, or menstrual cycle phase. This resilience may reflect the adaptive role of statistical learning, a mechanism that supports the formation of stable representations of environmental structure from early in development and across species (Santolin et al., 2020). In humans, both probability-based and serial order regularities can be learned and maintained for over a year, regardless of age (Tóth-Fáber et al., 2021). At the same time, differences in general performance, such as RT or accuracy, may reflect cultural factors, including unequal exposure to specific tasks (Dorfberger et al., 2009; Landauer et al., 1980), or developmental influences that shape procedural learning (Dorfberger et al., 2009). Altogether, these considerations suggest that performance differences are shaped by a complex interplay of biological, cultural, and experiential factors, whereas statistical learning appears to be a fundamental and resilient building block of human cognition.

## Ethics approval and consent to participate

⌷ The study was approved by the Research Ethics Committee of Eötvös Loránd University, Budapest, Hungary (protocol number: 2021/504).
⌷ All participants provided informed consent prior to enrollment.
⌷ The study was conducted in accordance with the principles of the Declaration of Helsinki.

## Availability of data and material

⌷ All behavioral data and the R code required to reproduce the study’s results are publicly available on the Open Science Framework (OSF) repository at https://osf.io/u82w9/overview.

## Authors’ contributions

Conceptualization: I. B., T. V., D. N.

Data curation: B. B., T. V.

Formal analysis: I. B., A. H., T. V.

Funding acquisition: T. V., D. N.

Investigation: B. B., T. V.

Methodology: I. B., B. B., A. H., T. V., D. N.

Project administration: T. V.

Resources: B. B., T. V.

Software: B. B., T. V.

Supervision: T. V., D. N.

Validation: I. B., T. V.

Visualization: I. B., T. V.

Writing – original draft: I. B., T. V., D. N.

Writing – review & editing: I. B., T. V., D. N.

