## Supplementary Materials for "No gender differences in predictive processing"

**Supplementary Methods**

***Participants***

In addition to the analyses based on gender, comparisons were conducted within the group of women participants. From the initial group of 334 women, 328 provided information about hormonal contraceptive use, 70 reported using hormonal contraceptives (M_age_ = 22.0 years, SD_Age_ = 3.46). For the purpose of comparison, a group of 70 women who did not use contraceptives was selected from the remaining 258 non-users (M_age_ = 22.0 years, SD_Age_ = 3.52). These two groups were matched as closely as possible by age using nearest-neighbor matching (1:1 ratio) implemented with the *matchit()* function from the *MatchIt* package in R.

Furthermore, a more detailed analysis was performed within the subgroup of women who were not using hormonal contraceptives. Out of the initial 334 women, 251 provided information about their menstrual cycle phase and confirmed that they were not taking hormonal contraceptives; only these participants were included in this analysis. It should be noted that answering these questions was not mandatory. The 251 women (M_Age_ = 22.53, SD_Age_ = 5.54) were categorized according to their self-reported menstrual cycle phase. The distribution was as follows: early follicular (n = 47), mid/late follicular (n = 60), ovulatory (n = 30), early luteal (n = 39), late luteal (n = 32), premenstrual (n = 17), and irregular (n = 26).

***Statistical analysis***

In the second set of analyses, focusing exclusively on women participants (n = 70 per group), hormonal contraceptive use (users vs. non-users) was included as the between-subjects factor, with non-users selected to be age-matched to the group of users using nearest-neighbor matching. Finally, in the third set of analyses, only women not using hormonal contraceptives were considered (n = 251), and menstrual cycle phase (early follicular, mid/late follicular, ovulatory, early luteal, late luteal, premenstrual, irregular) (Souza et al., 2012) was entered as a between-subjects factor.

For both sets of analyses, trial-level reaction times and accuracies were analyzed using Generalized Linear Mixed Models (GLMMs). When significant interactions were observed, post hoc pairwise comparisons were performed using estimated marginal means (emmeans package in R). Different adjustments for multiple testing were applied depending on the analysis. For the hormonal contraceptive analyses, Holm-adjusted comparisons were used for follow-up tests of significant interactions. For the menstrual-cycle analyses, Tukey’s HSD adjustment was used for all pairwise comparisons across menstrual cycle phases, as it provides better control of the family-wise error rate when all group pairs are compared (Kim, 2015).

**Supplementary Results**

***No evidence that hormonal contraceptive use influences visuomotor or statistical learning in reaction time measures***

To examine whether hormonal contraceptive use modulates predictive processing, a generalized linear mixed-effects model (GLMM) was fitted to trial-level reaction times (RTs). The model revealed a robust main effect of epoch (χ²(2) = 35.20, *p* < .001), indicating general skill learning throughout the task (Figure S1A). A significant main effect of triplet type (χ²(1) = 35.55, *p* < .001) confirmed reliable statistical learning, as participants responded faster to high-probability than to low-probability triplets (Figure S1C).

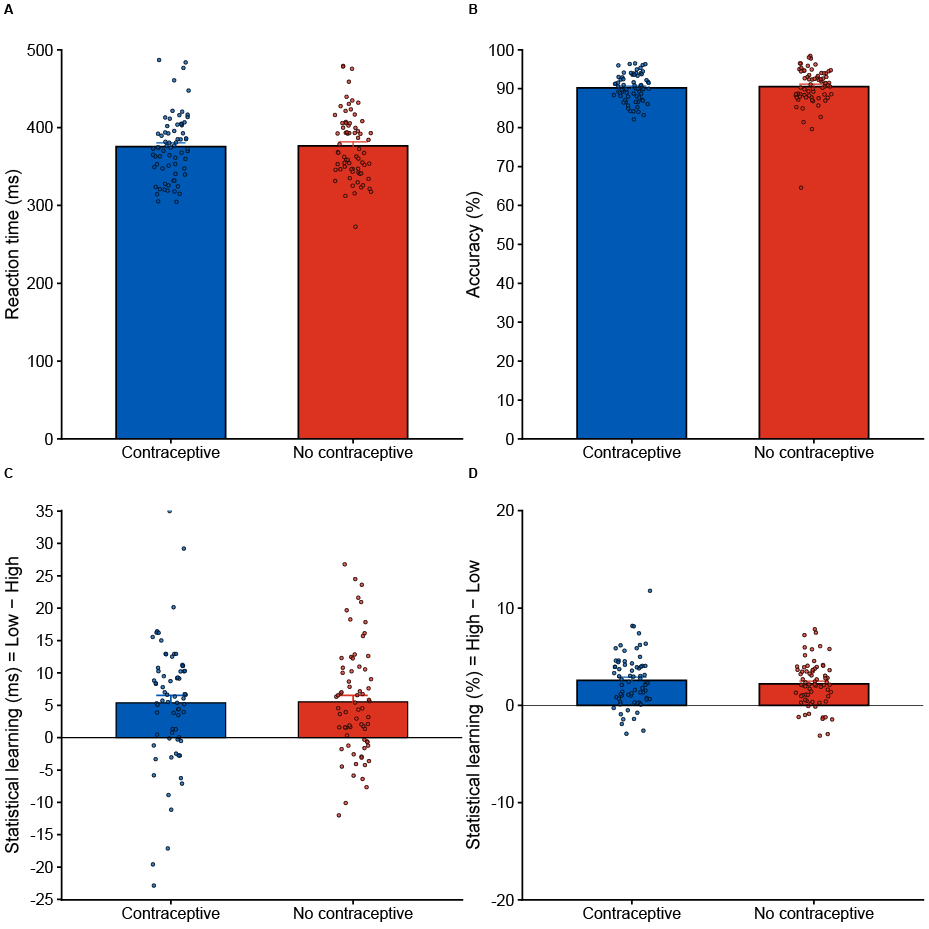

**Figure S1. Hormonal contraceptive use was not associated with differences in visuomotor or statistical learning.** Women using hormonal contraceptives are represented in blue and women not using hormonal contraceptives are represented in red throughout all panels. Bars represent group means and individual dots represent participant-level scores (n = 70 per group). The x-axis shows the two groups; the y-axis differs by panel. **A.** Baseline reaction time (RT; ms), computed as the average median RT across Epochs 1–3. Overall response speed was comparable between women using and not using hormonal contraceptives, consistent with the absence of a group effect in the RT GLMM. **B.** Baseline accuracy (%), computed as the average accuracy across Epochs 1–3. Accuracy levels were highly similar in the two groups, indicating no overall effect of contraceptive use on task performance. **C.** Statistical learning in RT, computed as Low − High (ms) and averaged across Epochs 1–3. Positive values indicate faster responses to high-probability than to low-probability triplets. Statistical learning was evident in both groups and did not differ between contraceptive users and non-users. **D.** Statistical learning in accuracy, computed as High − Low (%) and averaged across Epochs 1–3. Positive values indicate greater accuracy for high-probability than for low-probability triplets. Statistical learning was present in both groups and showed no meaningful differences as a function of hormonal contraceptive use.

In addition, a significant epoch × triplet type interaction (χ²(2) = 21.48, *p* < .001) demonstrated that the magnitude of statistical learning changed across the course of the task. Crucially, hormonal contraceptive use was not associated with overall RT performance, as evidenced by the absence of a main effect of group (χ²(1) < 0.01, *p* = .991). More importantly, contraceptive use did not interact with triplet type (χ²(1) = 0.18, *p* = .670), indicating that statistical learning was comparable between women who used hormonal contraceptives and those who did not. Likewise, the three-way interaction between group, epoch, and triplet type was entirely absent (χ²(2) = 0.11, *p* = .944), demonstrating that the trajectory of statistical learning over time was statistically indistinguishable between the two groups (Figure S1C). Together, these findings provide no evidence that hormonal contraceptive use influences predictive processing as measured by RTs.

***No evidence that hormonal contraceptive use influences statistical learning in accuracy measures***

A GLMM was fitted to the accuracy data using the same fixed- and random-effects structure as in the RT analysis. The accuracy model confirmed robust visuomotor learning throughout the task, evidenced by a highly significant main effect of epoch (χ²(2) = 22.14, *p* < .001) (Figure S1B). Furthermore, statistical learning was clearly established, as the main effect of triplet type was highly significant (χ²(1) = 91.81, *p* < .001), indicating that participants responded more accurately to high-probability than to low-probability triplets. The model also revealed a significant interaction between epoch and triplet type (χ²(2) = 10.39, *p* = .006), demonstrating that the magnitude of statistical learning evolved across the epochs. Follow-up post-hoc comparisons revealed that the statistical learning effect was significantly smaller in Epoch 1 than in Epoch 2 (OR = 1.14, *p* = .011) and Epoch 3 (OR = 1.15, *p* = .010), whereas Epochs 2 and 3 did not differ (OR = 1.01, *p* = .893), suggesting that learning emerged early and subsequently stabilized. Although the interaction between group and epoch approached significance (χ²(2) = 5.65, *p* = .059), this effect did not reach the conventional significance threshold.

The GLMM provided no evidence for a main effect of hormonal contraceptive use (χ²(1) = 1.09, *p* = .297). Most importantly, the interaction between hormonal contraceptive use and triplet type was not significant (χ²(1) = 0.16, *p* = .686), indicating that women who used hormonal contraceptives did not differ from women who did not use hormonal contraceptives in their statistical learning. Additionally, the three-way interaction between group, epoch, and triplet type was completely absent (χ²(2) = 0.63, *p* = .731), confirming that the trajectory of statistical learning over time was comparable between the two groups (Figure S1D). Therefore, these findings provide no evidence for differences in predictive processing as a function of hormonal contraceptive use.

***No evidence that menstrual cycle phase influences statistical learning in reaction time measures***

To analyze the trial-by-trial dynamics of statistical learning across the menstrual cycle, a GLMM was fitted to the RT data. The RT model confirmed robust visuomotor learning throughout the task, evidenced by a highly significant main effect of epoch (χ²(2) = 52.03, *p* < .001). Furthermore, statistical learning was clearly established, as the main effect of triplet type was highly significant (χ²(1) = 61.81, *p* < .001), indicating that participants responded faster to high-probability than to low-probability triplets. The model also revealed a significant interaction between epoch and triplet type (χ²(2) = 30.31, *p* < .001), demonstrating that the magnitude of statistical learning evolved across the epochs.

Addressing the primary research question regarding menstrual-cycle differences in predictive processing, the GLMM provided no evidence for a main effect of menstrual cycle phase (χ²(6) = 9.90, *p* = .129). Most importantly, the interaction between menstrual cycle phase and triplet type was not significant (χ²(6) = 4.63, *p* = .592), indicating that women in different menstrual-cycle phases did not differ in their sensitivity to statistical regularities. Additionally, the three-way interaction between menstrual cycle phase, epoch, and triplet type was completely absent (χ²(12) = 5.43, *p* = .942), confirming that the trajectory of statistical learning over time was statistically indistinguishable across menstrual-cycle phases (Figure S2C).

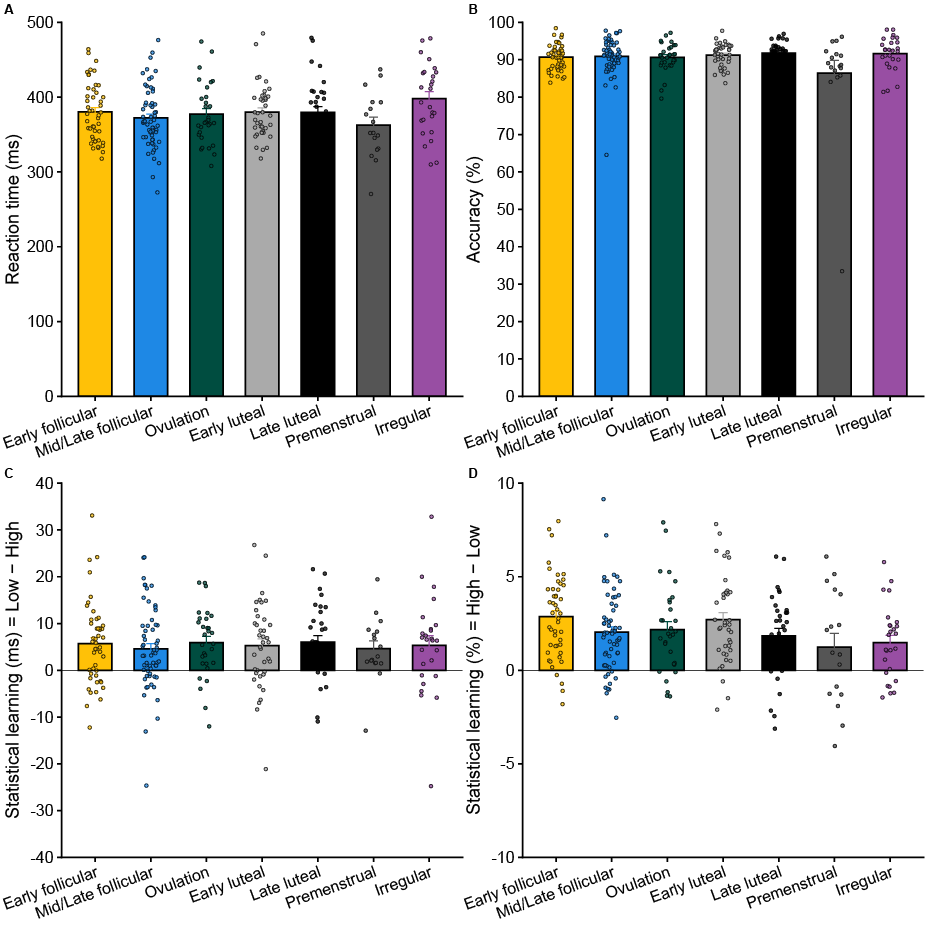

**Figure S2. Absence of menstrual cycle effects on visuomotor performance and statistical learning.** Panels show group means ± SEM overlaid with individual data points; the x-axis shows self-reported cycle phases (Early follicular, Mid/Late follicular, Ovulation, Early luteal, Late luteal, Premenstrual, Irregular). **A.** Reaction time (RT). Mean RTs (ms) across all trials. Overall response speed did not differ significantly across menstrual-cycle phases, consistent with the absence of a main effect of cycle phase in the RT GLMM. **B.** Accuracy. Mean accuracy (%) across all trials. Accuracy levels were generally comparable across menstrual-cycle phases. Although visual inspection suggests slightly lower accuracy in the premenstrual group, the overall effect of cycle phase did not reach significance and should be interpreted cautiously given the relatively small sample size of this subgroup. **C.** Statistical learning in RT. Statistical learning was computed as Low − High (ms), with positive values indicating faster responses to high-probability than to low-probability triplets. Statistical learning was robust across all menstrual-cycle phases and did not differ significantly as a function of cycle phase. **D.** Statistical learning in accuracy. Statistical learning was computed as High − Low (%), with positive values indicating greater accuracy for high-probability than for low-probability triplets. Although visual inspection suggests somewhat smaller learning effects in the premenstrual group, follow-up comparisons revealed no significant differences between menstrual-cycle phases after correction for multiple comparisons. Overall, statistical learning in accuracy was comparable across menstrual-cycle phases, providing no reliable evidence that menstrual-cycle phase modulates predictive processing.

***No evidence that menstrual cycle phase influences statistical learning in accuracy measures***

A GLMM was fitted to the accuracy data using the same fixed- and random-effects structure as in the RT analysis. The accuracy model confirmed robust general skill learning throughout the task, evidenced by a highly significant main effect of epoch (χ²(2) = 34.02, *p* < .001). Furthermore, statistical learning was clearly established, as the main effect of triplet type was highly significant (χ²(1) = 152.10, *p* < .001), indicating that participants responded more accurately to high-probability than to low-probability triplets. The model also revealed a significant interaction between epoch and triplet type (χ²(2) = 12.92, *p* = .002), demonstrating that the magnitude of statistical learning evolved across the epochs. Follow-up comparisons revealed that the statistical-learning effect was significantly larger in Epoch 3 than in Epoch 1 (OR = 1.15, *p* = .001) and Epoch 2 (OR = 1.09, p = .047), whereas Epochs 1 and 2 did not differ significantly (OR = 1.05, *p* = .179), indicating a gradual increase in statistical learning across the task.

Addressing the primary research question regarding menstrual-cycle differences in predictive processing, the GLMM revealed no significant main effect of menstrual cycle phase (χ²(6) = 10.88, *p* = .092). The interaction between menstrual cycle phase and triplet type reached trend-level significance (χ²(6) = 11.40, *p* = .077). Additionally, the three-way interaction between menstrual cycle phase, epoch, and triplet type was not significant (χ²(12) = 11.17, *p* = .515), confirming that the trajectory of statistical learning over time was comparable across menstrual-cycle phases (Figure S2D). Taken together, these findings suggest that the trend-level interaction is unlikely to reflect a reliable phase-specific modulation of predictive processing.

**Supplementary Figures**

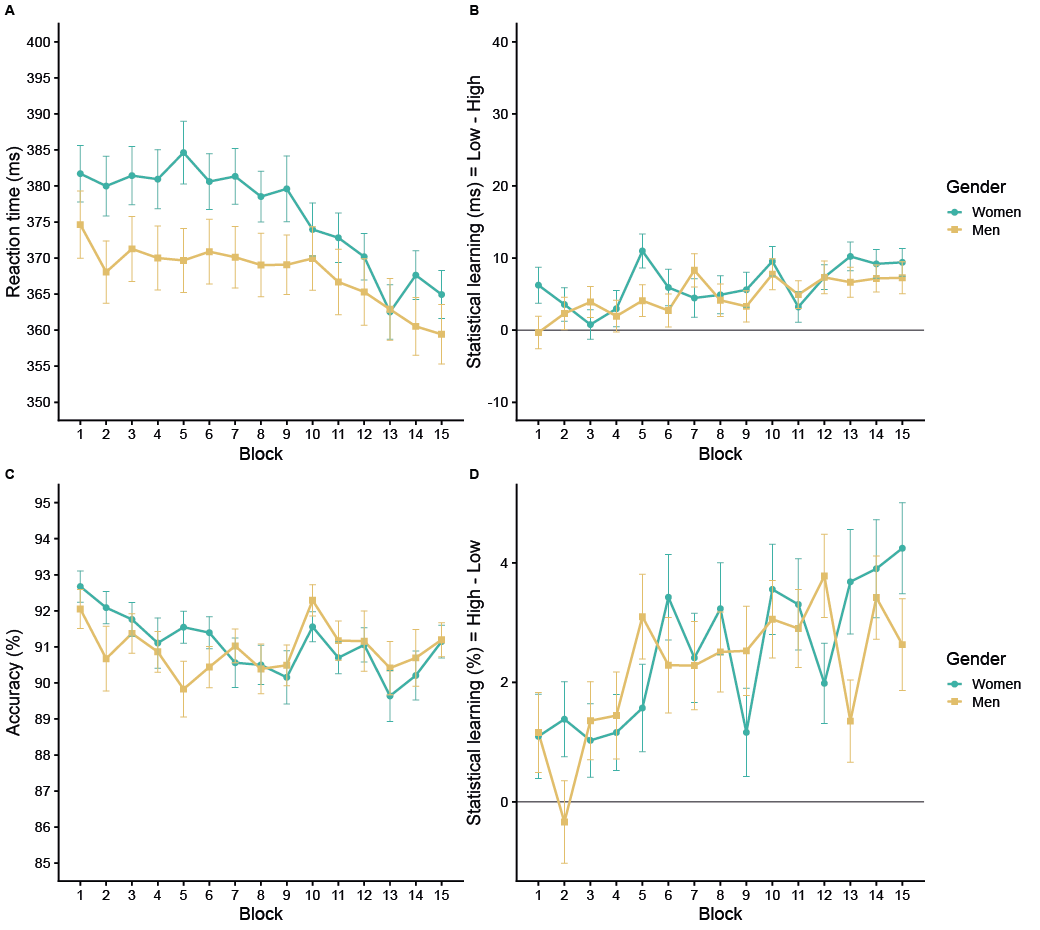

**Figure S3. Block-by-block trajectories of visuomotor performance and statistical learning by gender.** Women are represented in green and men in yellow throughout all panels. Lines represent group means and error bars indicate the standard error of the mean (SEM). The x-axis shows the 15 blocks of the ASRT task. **A.** Reaction times (RTs; ms) across blocks. RTs gradually decreased throughout the task, indicating general skill learning. Women exhibited generally slower RTs than men; however, the pattern of improvement across blocks was highly similar between groups. B. Statistical learning in RT, computed as Low − High (ms) for each block. Positive values indicate faster responses to high-probability than to low-probability triplets. Statistical learning was evident across the task and showed a gradual increase over blocks, with highly similar trajectories in women and men. **C.** Accuracy (%) across blocks. Accuracy remained relatively stable throughout the task, with comparable performance in women and men. **D.** Statistical learning in accuracy, computed as High − Low (%) for each block. Positive values indicate greater accuracy for high-probability than for low-probability triplets. Statistical learning was present across blocks and increased modestly over time, with no clear evidence of gender-related differences in learning trajectories.
